# Absence of heme at the catalytic site of heme oxygenase-2 triggers its lysosomal degradation

**DOI:** 10.1101/2020.06.21.163881

**Authors:** Liu Liu, Arti B. Dumbrepatil, Angela S. Fleischhacker, E. Neil G. Marsh, Stephen W. Ragsdale

## Abstract

Heme oxygenase-2 (HO2) and −1 (HO1) catalyze heme degradation to biliverdin, CO, and iron, forming an essential link in the heme metabolism network. Tight regulation of the cellular levels and catalytic activities of HO1 and HO2 is important for maintaining heme homeostasis. While transcriptional control of HO1 expression has been well-studied, how the cellular levels and activity of HO2 are regulated remains unclear. Here, the mechanism of post-translational regulation of cellular HO2 level by heme is elucidated. Under heme deficient conditions, HO2 is destabilized and targeted for degradation. In HO2, three heme binding sites are potential targets of heme-dependent regulation: one at its catalytic site; the others at its two heme regulatory motifs (HRMs). We report that, in contrast to other HRM-containing proteins, the cellular protein level and degradation rate of HO2 are independent of heme binding to the HRMs. Rather, under heme deficiency, loss of heme binding to the catalytic site destabilizes HO2. Consistently, a HO2 catalytic site variant that is unable to bind heme exhibits a constant low protein level and an enhanced protein degradation rate compared to the wild-type HO2. However, cellular heme overload does not affect HO2 stability. Finally, HO2 is degraded by the lysosome through chaperone-mediated autophagy, distinct from other HRM-containing proteins and HO1, which are degraded by the proteasome. These results reveal a novel aspect of HO2 regulation and deepen our understanding of HO2’s role in maintaining heme homeostasis, paving the way for future investigation into HO2’s pathophysiological role in heme deficiency response.

---

Heme is a commonly used prosthetic group in a myriad of biological processes. High levels (>1 μM) of free heme are seen in many pathological processes and diseases, such as cardiovascular diseases (CVD) (1). The cytotoxicity of heme arises from the overproduction and accumulation of reactive oxygen species (ROS), leading to oxidative stress and tissue damage (2,3). Heme oxygenase (HO) is the only known mammalian enzyme that catalyzes the degradation of heme, thus maintaining systemic heme homeostasis. Furthermore, HO converts heme to biliverdin (further reduced to bilirubin) and CO, which play essential roles in cytoprotection against oxidative stress (4-7). Therefore, understanding the regulation of HO activity is important for human health.

In mammals, two major isoforms of HO have been identified: HO1 and HO2. These two enzymes share high sequence homology and have very similar structures (4,5,8), especially for regions around the catalytic site where heme binds and is metabolized. The catalytic site binds heme with high affinity through a conserved histidine (His45 in HO2, His25 in HO1) as the proximal ligand and an aquo (H_x_O) ligand H-bonded to Gly159 (Gly139 in HO1) as the distal ligand (Fig. 1A). Consistent with their high homology, these enzymes possess very similar heme degradation activities (8).

**Figure 1.**
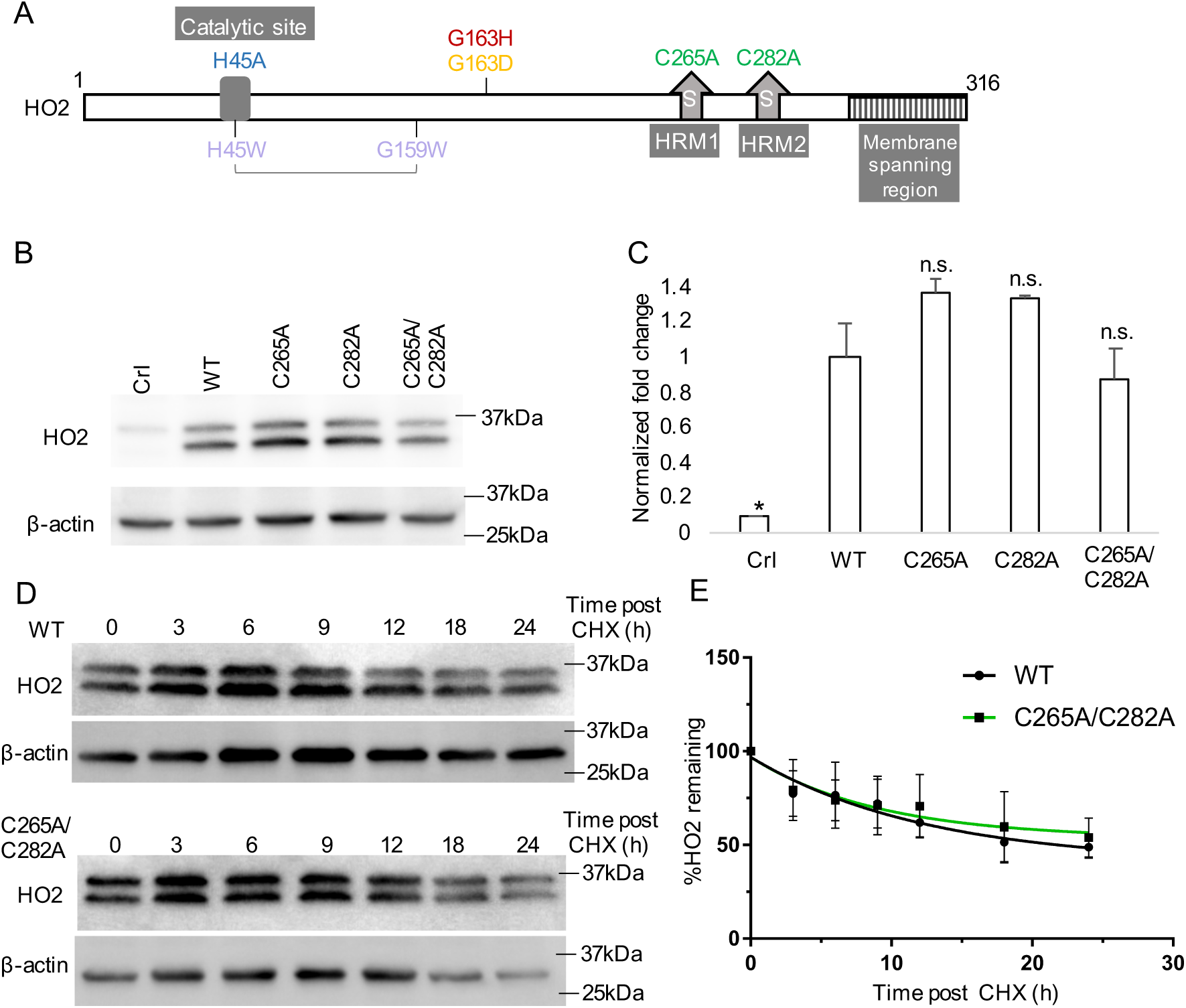
Western blotting analysis indicates that lack of heme binding to HRMs does not affect HO2 protein stability and degradation. (***A***) A diagram of HO2 sequence with all variants used in this article indicated as follows: HRM mutations (*green*); Catalytic site mutations: loss of heme binding (*light purple*), weaker heme binding (*blue and yellow*) and tighter heme binding mutations (*red*). (***B****) A* representative western blot demonstrating steady-state expression levels of HO2 in HEK293 cells transiently transfected with transfection reagent only (*Crl*, control), or the corresponding expression vectors for wild-type HO2 (*WT*) and HRM mutants (*C265A, C282A or C265A/C282A*) for 24h. Samples were probed with anti-HO2 antibody (*top panel*), or anti-actin antibody as a loading control (*bottom panel*). (***C***) The intensities of the bands in (*B*) were quantified by densitometry, and the mean intensities ± S.D. of at least three independent experiments were plotted. *p<0.05; n.s. p>0.05 vs WT. (***D***) HEK293 cells transfected with WT-HO2 (*top experiment*) or the C265A/C282A mutant (*bottom experiment*) were treated with CHX (35 μg/ml) for indicated time before collection. Cell lysates were probed by western blot with anti-HO2 antibody or anti-actin antibody as loading control. (***E***) The intensities of the bands in (*D*) were quantified by densitometry, and the mean intensities ± S.D. of three independent experiments were plotted. Data were fitted into a one-phase decay curve using GraphPad Prism 8 to calculate protein half-lives for WT-HO2 (*black*) and C265A/C282A mutant (*green*), respectively.

Despite these similarities, HO1 exists in a tissue-specific and inducible manner, with its expression stimulated at the transcriptional level by various conditions, such as infection, excess heme, and oxidative stress (4). In contrast, HO2 is constitutively expressed and highly concentrated in brain, testes and neural tissues (4). HO2 is considered to provide the baseline heme detoxification activity under normal conditions and to be cytoprotective under stressed conditions, when or where HO1 is not available, such as ischemic stroke (9,10). As of now, little is known about how HO2 is regulated to maintain its proper heme detoxification and cytoprotective functions.

Notably, the biggest difference between those two enzymes exists in the region C-terminal to the catalytic core, where HO1 harbors a PEST domain that is usually associated with proteasomal degradation of short-lived proteins (11,12). HO2, however, possesses two heme regulatory motifs (HRMs) in this corresponding region (Fig. 1A). The HRM is a conserved sequence, featured by a Cys-Pro dipeptide core with Cys serving as the heme ligand (13). In HO2, there are two HRMs, centered at Cys265 and Cys282, each of which binds heme under reducing conditions; yet in an oxidizing environment, they form a disulfide bond and are unable to bind heme. Previous *in cellulo* and *in vitro* studies showed that this thiol-disulfide redox switch is 60-70% reduced under normal physiological conditions and exhibits a midpoint potential of −200 mV, which is within the range of ambient cellular redox potential (14). Moreover, the binding affinities for heme of the two HRMs match well with the cellular regulatory heme pool level (14-17). Thus, this redox-gated heme binding appears to be physiologically relevant.

Several key proteins within the heme metabolism network are subject to HRM-mediated heme-dependent regulation of their expression, degradation, and activity (18). For instance, heme binding to the HRMs of Bach1, a transcriptional repressor of the HO1 gene, disrupts its DNA binding activity (19) and triggers the ubiquitination and proteasomal degradation of Bach1 protein, leading to HO1 induction (20). In another case, the cellular level of δ-aminolevulinic acid synthase 1 (ALAS1), the rate-limiting enzyme in heme biosynthesis, is also mediated by heme-HRM interaction through a negative feedback mechanism (21). A recent survey of HRM-containing proteins demonstrated that the most common HRM function is to promote protein degradation upon heme binding (18). It has been speculated that heme binding to the HRM may serve as a molecular signature for recognition by an E3 ubiquitin ligase so as to target those HRM-containing proteins for proteasomal degradation (20,22). Therefore, we hypothesized that heme binding to the two HRMs in HO2 modulates HO2 protein degradation.

In order to understand how heme regulates HO2 protein stability and degradation, we constructed HO2 mutants with various heme binding abilities at each of the three heme binding sites (Fig. 1A). Our work shows that, unlike other HRM-containing proteins, the HRMs in HO2 do not mediate protein degradation. In addition, HO2 is degraded by the lysosome through chaperone-mediated autophagy instead of the ubiquitin-proteasome pathway used for other HRM-containing proteins (18) or the endoplasmic reticulum (ER)-associated protein degradation (ERAD) pathway used for HO1 (12). Rather, we find an unexpected regulatory function of the catalytic site of HO2 beyond its enzymatic activity: the cellular protein level of HO2 is controlled by heme occupancy at the catalytic site. Under the condition of heme deficiency, heme is absent from HO2’s catalytic site, resulting in a lower protein level. This low protein level phenomenon is well mimicked by a HO2 variant that cannot accommodate heme at the catalytic site by default. Our study reveals important insight into how this housekeeping HO is regulated to maintain heme homeostasis and adds one potential link in cellular response against heme deficiency.

## RESULTS

### HO2 stability and degradation is independent of heme binding to the HRMs

Given that the most common role of HRMs is to promote protein degradation upon binding heme, we first asked whether the two HRMs in HO2 also control protein stability and degradation. Earlier, we showed that substituting Cys265 or Cys282 with Ala abolishes heme binding to HRM1 or HRM2, respectively (15,16). Therefore, we introduced mutations, i.e. C265A, C282A and C265A/C282A, in the HRMs of HO2 to disrupt heme binding, anticipating that the mutations would lead to HO2 stabilization. The three variants and wild-type HO2 were constructed in a mammalian expression vector to dissect their properties under a cellular environment. HEK293 cells were transfected to overexpress the wild-type and variants of HO2 and their steady-state expression levels were studied. When probing HO2 protein levels with a rabbit-polyclonal anti-HO2 antibody, we observed two bands both corresponding to HO2 in cell extracts, with the higher molecular weight one assigned to full-length HO2 (Fig. 1B). The lower molecular weight band has been often observed with high cellular HO2 levels (11,14,23). Since both bands correspond to HO2, they both have been taken into account when quantifying their intensities.

Extracts from non-transfected cells yielded very low levels of endogenous protein (Fig. 1B, Crl lane). In contrast to what has been observed with ALAS1 or Bach1, in which a loss of function mutation in the HRM leads to a higher cellular level of protein (21), all three HRM variants of HO2 maintained a steady-state expression level very similar to that of wild-type protein (Fig. 1B-1C). Furthermore, we compared the half-life of the C265A/C282A variant to that of wild-type protein using a cycloheximide (CHX) chase assay (24). In cells treated with CHX to repress protein synthesis, the HO2 protein level in total cell extracts decreased exponentially with increasing chase time (Fig. 1D-1E), whereas in the control group treated with vehicle DMSO, HO2 protein levels were stable (Fig. S1). Fitting the data to a one phase decay equation yields very similar degradation rates of the C265A/C282A variant and wild-type protein (Fig. 1E). These results suggest that heme binding to the HRMs does not significantly impact the stability or degradation rate of HO2.

### HO2 protein level decreases in response to lowered cellular heme concentration

We next examined how HO2 protein level responds to changes in cellular heme concentration. We first quantified total cellular heme in HEK293 cells grown in a basal culture medium. With single cell volume estimated as 3×10^−12^ L (25), the cellular total heme concentration was calculated to be 3.16 ± 0.03 μM. However, the regulatory heme pool, which has been reported to be around 20-340 nM (17,26,27), is far lower than total heme. Considering the affinity of the HRMs for heme (15), as well as the limited availability of the cellular regulatory heme pool (17), the HRMs may not be fully heme replete under regular culture conditions.

Addition of the heme biogenesis inhibitor succinylacetone (SA) to the heme-depleted (HD) medium for 72 h depletes the regulatory heme pool (17) and decreases total cellular heme by ∼25% (Fig. 2A). Supplementing with hemin after SA treatment replenishes cellular heme, and depending on the dose of hemin added, the cellular heme increases by up to 3-fold, (Fig. 2A). While the degradation of other HRM-containing proteins, such as ALAS1 or Bach1 are accelerated by hemin treatment (20,21), we did not observe any effect of increasing the cellular heme content on HO2 expression (Fig. 2B). This result reinforces our conclusion that heme binding to the HRMs of HO2 does not regulate protein stability or degradation. Quite surprisingly, however, we observed a decrease in HO2 steady-state protein level in response to SA treatment, indicating that HO2 protein is destabilized under heme deficient condition (Fig. 2B-2C). Therefore, we set out to understand how heme levels influence HO2 stability and postulated that lack of heme binding to the catalytic site, but not the HRMs, may be responsible for the observed destabilization of HO2.

**Figure 2.**
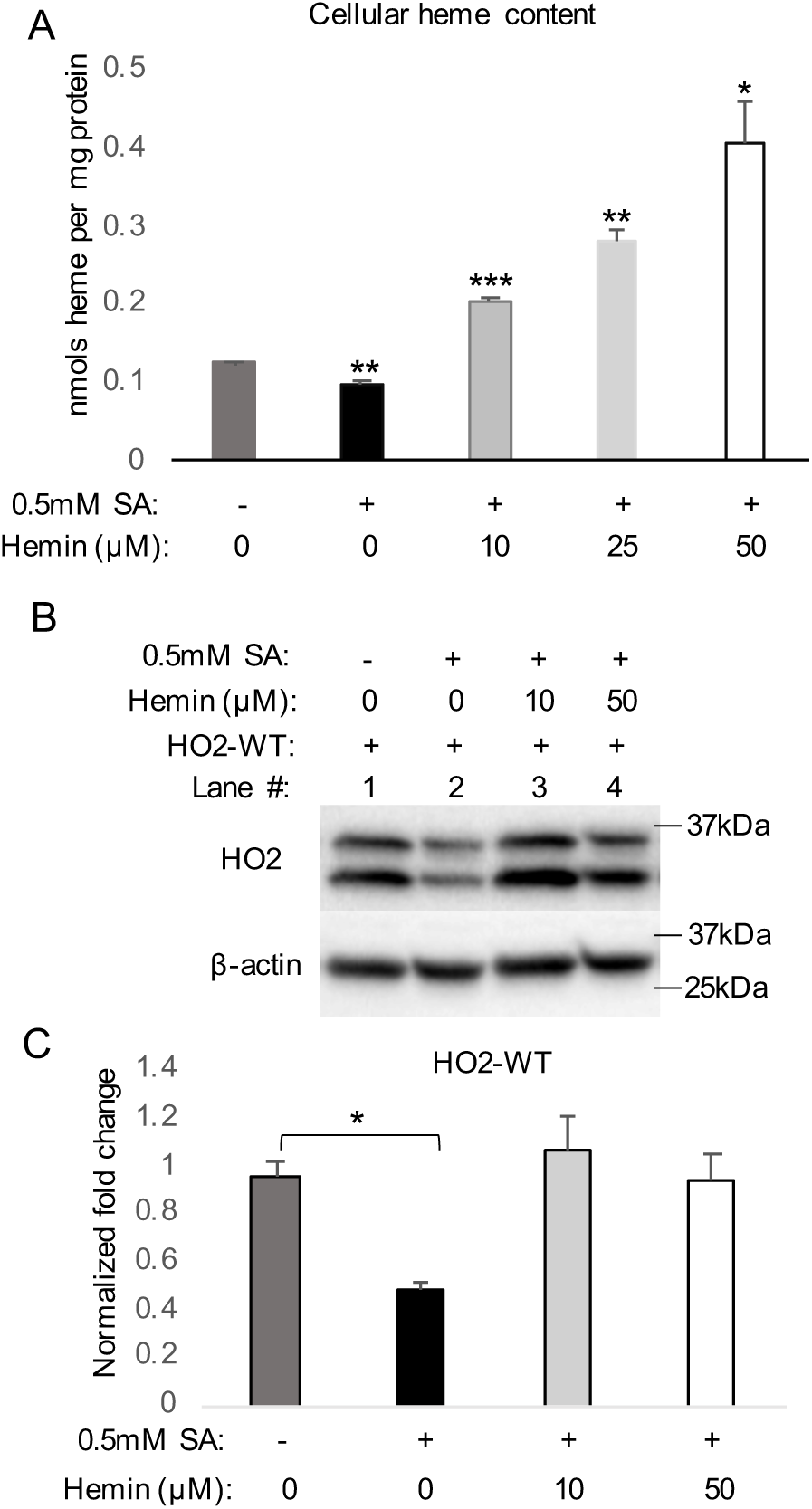
The degradation of HO2 is enhanced in response to lowered cellular heme. (***A***) Oxalic acid assay quantification of cellular heme concentration of HEK293 cells cultured in heme-depleted (HD) media with different treatment conditions, n=3. (***B***) HEK293 cells cultured in heme-depleted (HD) media were transfected to express wild-type HO2 (WT) and treated with 0.5 mM SA for 48h. Then 0, 10 or 50 μM of hemin was supplied to the cultures as indicated for 24h before sample collection. Samples were probed with anti-HO2 antibody (*top panel*), or anti-actin antibody as a loading control (*bottom panel*). (***C***) The intensities of the bands in (*B*) were quantified by densitometry, and the mean intensities ± S.D. of two independent experiments were plotted. For (*A*) and (*C*), *p<0.05; **p<0.01 and ***p<0.001 vs condition in lane 1.

### Lack of heme occupancy at the catalytic site leads to HO2 degradation

Due to its high affinity for heme (K_d_ ∼3.6 nM) (28), in a regular cellular environment where the regulatory heme pool is around 20-340 nM (29), the catalytic site of wild-type HO2 would most likely be heme-replete. To explore the role of heme binding to the catalytic site in regulating HO2 stability, we generated a catalytic site loss-of-function mutant, H45W/G159W, for expression in HEK293 cells. *In vitro* experiments have confirmed that in H45W/G159W variant, there is no detectible heme binding at the catalytic site and that the overall structure and heme binding to the HRMs are unperturbed (30). As expected, no changes were observed in steady-state expression levels of the H45W/G159W mutant upon cellular heme level variation (Fig. 3A-3B). Rather, it exhibited a consistently lower protein level when compared with its wild-type counterpart (Fig. 3E, Fig. S1A).

**Figure 3.**
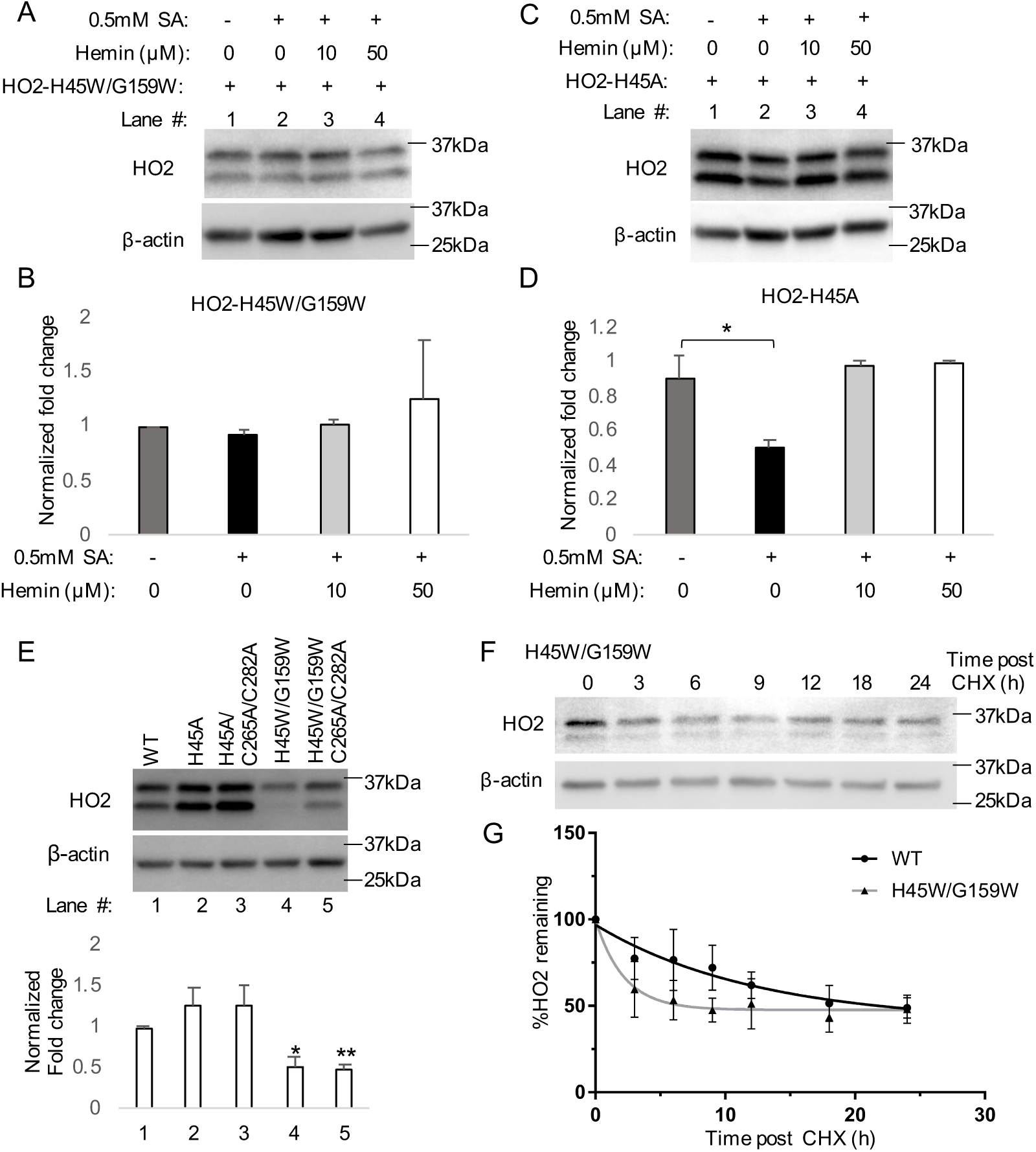
Binding of heme to the catalytic site stabilizes HO2 protein. (***A, C***) HEK293 cells cultured in HD media were transfected with HO2-H45W/G159W *(A)* or H45A *(C)*, then treated with 0.5 mM SA for 48h. 0, 10 or 50 μM of hemin was supplied to the cultures as indicated for 24h before sample collection. Representative western blots for samples probed with anti-HO2 antibody (*top panel*), or anti-actin antibody as a loading control (*bottom panel*) are shown. (***B, D***), The intensities of the bands in *(A)* and *(C)* were quantified respectively by densitometry, and the mean intensities ± S.D. of at least three independent experiments were plotted. *p<0.05 compared with lane 1. (***E***) A representative western blot demonstrating steady-state expression levels of HO2 in HEK293 cells transiently transfected with the corresponding expression vector for wild-type HO2 (*WT*), catalytic site mutants (*H45A or H45W/G159W*) or catalytic site mutants combined with HRM mutations (*H45A/C265A/C282A* or *H45W/G159W/C265A/C282A*). Samples were probed with anti-HO2 antibody (*top panel*), or anti-actin antibody as a loading control (*bottom panel*). The intensities of the bands were quantified by densitometry, and the mean intensities ± S.D. of at least three independent experiments were plotted. *p<0.05; **p<0.01 vs lane 1. (***F)*** HEK293 cells transfected with H45W/G159W variant were treated with CHX (35 μg/ml) for indicated time before collection. Cell lysates were probed by western blot with anti-HO2 antibody (*top panel*), or anti-actin antibody as a loading control (*bottom panel*). A representative western blot for wild-type HO2 degradation has been shown in **Figure. 1**. (***G***) The intensities of the bands in (F) were quantified by densitometry, and the mean intensities ± S.D. of three independent experiments were plotted. Data were fitted into a one-phase decay curve using GraphPad Prisma 8 to calculate protein half-life for H45W/G159W (*light gray*), which is compared with WT-HO2 (*black*).

Because the H45W/G159W variant cannot bind heme, it is also necessarily catalytically inactive. Therefore, to dissect whether the enzymatic activity contributes to the regulation of HO2 stability, we introduced another HO2 variant, H45A, in which the catalytic core still accommodates heme but lacks catalytic activity (Fig. S2, Table S1) (31). Similar to the wild-type protein, the H45A variant showed a decreased steady-state expression level when grown in the presence of SA, which lowers cellular heme level (Fig. 3C-3D). This result suggests that it is the heme occupancy at the catalytic site, not enzymatic activity, that controls HO2 stability.

To further validate that the heme-dependent HO2 destabilization is contributed by lack of heme binding to the catalytic site only, we probed steady-state HO2 levels when HRMs were present or absent (fulfilled with C265A/C282A) in H45A and H45W/G159W mutants. A decrease in steady-state HO2 protein level was only observed when the catalytic core was in the “apo” state, i.e., harboring H45W/G159W mutations, regardless of whether the HRMs were present or absent (Fig. 3E). To ensure the observed lower steady-state level of the H45W/G159W variant indeed results from an enhanced protein degradation, its protein half-life was measured. We found the half-life of the H45W/G159W variant (1.5 h) to be ∼6-fold shorter than that of wild-type HO2 (8.6 h) (Fig. 3F-3G). To better characterize this fast-degraded variant, we collected more data time points between 0 h and 3 h post-CHX treatment and observed a 12-fold decrease in the half-life (∼0.7 h) of the H45W/G159W variant (Fig. S3).

Since the H45A variant lacks a strong heme iron axial ligand, the majority of the heme accommodated by the H45A variant is only weakly associated with the protein (Fig. S2). To better distinguish the effect of heme binding versus heme metabolism on HO2 stability, we considered other variants in which heme affinity to the catalytic site is not as markedly decreased as H45A. We measured heme off-rates of these variants; heme off-rates are a reflection of heme-binding affinity (*K*_*d*_) since it has been shown by our group and others that heme on-rates of many hemoproteins, despite structural differences, are very similar (15,28,32). The G163D variant is catalytically inactive (Fig. S2 and Table S1) but, unlike the H45A variant, exhibits a very similar affinity for heme at the catalytic site as wild-type HO2 (Table S1). The marked effect of heme occupancy at the catalytic site on HO2 stability also led us to explore whether increasing heme affinity of the catalytic site would further stabilize HO2 protein, leading to a higher steady-state HO2 level than wild-type. Therefore, the G163H variant, also catalytically inactive (Table S1), however with a 100-fold higher heme affinity of the catalytic site (30), was tested. Surprisingly, both the G163D and G163H variants showed similar steady-state protein levels as wild type, and the G163H variant has a similar half-life as wild-type HO2 (Fig. 4A-4D). Our results thus suggest that the wild-type protein, and even the H45A variant, already is sufficiently heme-replete in a regular cellular environment, so that no further stabilization can be achieved by increasing the heme affinity of the catalytic site. We postulate that the only time we would be able to observe a decrease in protein stability with G163H is when the regulatory heme pool dropped into the pM level.

**Figure 4.**
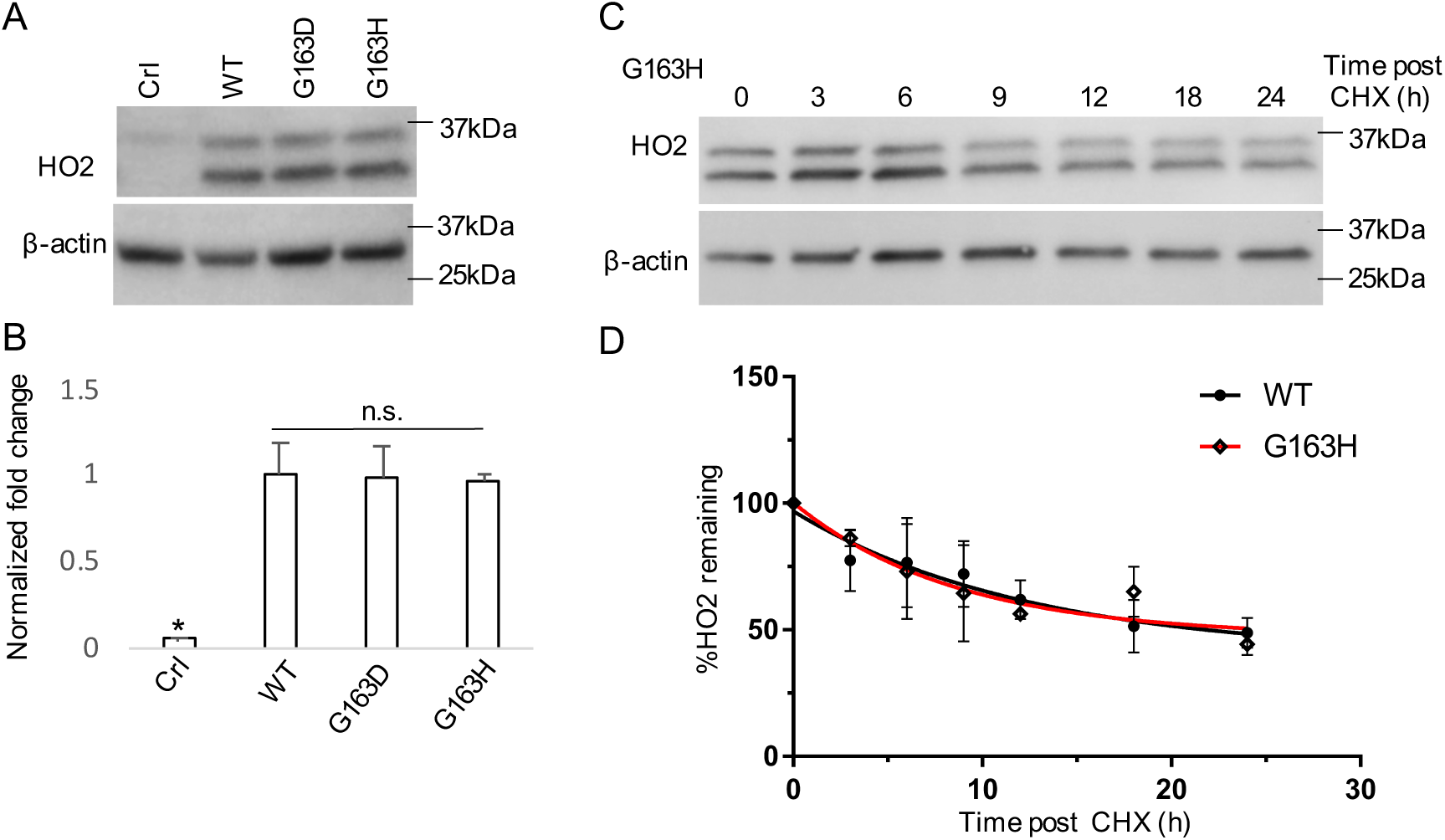
A tighter heme binding to the catalytic site does not further stabilize HO2 protein. (***A***) A representative western blot demonstrating steady-state expression levels of HO2 in HEK293 cells transiently transfected with transfection reagent only (Crl, control), or the corresponding expression vectors for wild-type HO2 (*WT*) and catalytic core mutants (G163D or G163H). Samples were probed with anti-HO2 antibody (*top panel*), or anti-actin antibody as a loading control (*bottom panel*). (***B***) The intensities of the bands in (*A*) were quantified by densitometry, and the mean intensities ± S.D. of at least three independent experiments were plotted. * p<0.05 vs WT; n.s. p>0.05 vs WT. (***C***) HEK293 cells transfected with the tighter binding mutant G163H were treated with CHX (35 μg/ml) for indicated time before collection. Cell lysates were probed by western blot with anti-HO2 antibody (*top panel*), or anti-actin antibody as a loading control (*bottom panel*). A representative western blot for wild-type HO2 degradation has been shown in **Figure 1**. (***D***) The intensities of the bands in (*C*) were quantified by densitometry, and the mean intensities ± S.D. of three independent experiments were plotted. Data were fitted into a one-phase decay curve using GraphPad Prism 8 to calculate protein half-life for G163H (*red*), which is compared with WT-HO2 (*black*).

### Immunofluorescence microscopy reveals heme binding to the catalytic site or the HRMs does not affect HO2 subcellular localization

Both HO isoforms were reported to be residents of the ER as type II membrane proteins, anchored by their single C-terminal hydrophobic helix (11,33). Because heme markedly affects HO2 stability, it is important to examine whether the introduced mutations or heme binding to the catalytic site or the HRMs altered HO2 subcellular localization. We attempted to visualize endogenous HO2; however, immunostaining using anti-HO2 antibody was insufficiently sensitive to detect HO2 when using the same exposure as for the overexpressed protein (Fig. 5, panel 1). When using higher exposure, we observed foci among whole cells which likely reflects non-specific binding of the antibody (Fig. S4). Therefore, we used anti-FLAG antibodies to visualize FLAG-tagged HO2 expressed in HEK293 cells. Calnexin, an ER-integrated chaperone that assists in the proper folding and quality control of glycoproteins, was used as the ER marker (34,35), while nuclei were shown by DAPI staining. To ensure that the N-terminal FLAG tag does not alter HO2 properties, we analyzed the steady-state expression levels of a FLAG-tagged version of wild-type HO2 and the two heme binding site mutants, C265A/C282A (HRMs) and H45W/G159W (catalytic site). Their expression levels matched those of untagged HO2 (Fig. S5A-S5B). Notably, only one band is observed by immunoblotting with anti-HO2 antibodies and it corresponds to full-length HO2. We conclude that expression of the untagged-and FLAG-tagged HO2 in different expression vectors (pcDNA3.1 and pQCXIP, respectively) affects how HO2 is processed in the cells. Indeed, the FLAG-tag itself does not affect HO2 properties, because we observe the same expression pattern for FLAG-tagged and untagged-HO2 in the pcDNA3.1 vector (Fig. S5C-S5D). Wild-type HO2 co-localized with calnexin and also exhibited a clear pattern surrounding the nuclei (Fig. 5, panel 2). Loss of heme binding at either the HRMs or the catalytic site did not alter HO2’s subcellular localization, suggesting that association of HO2 with the ER is heme-independent (Fig. 5, panel 3,4). The results also validate that the mutations introduced in the catalytic site and HRMs did not alter HO2’s association with the ER membrane.

**Figure 5.**
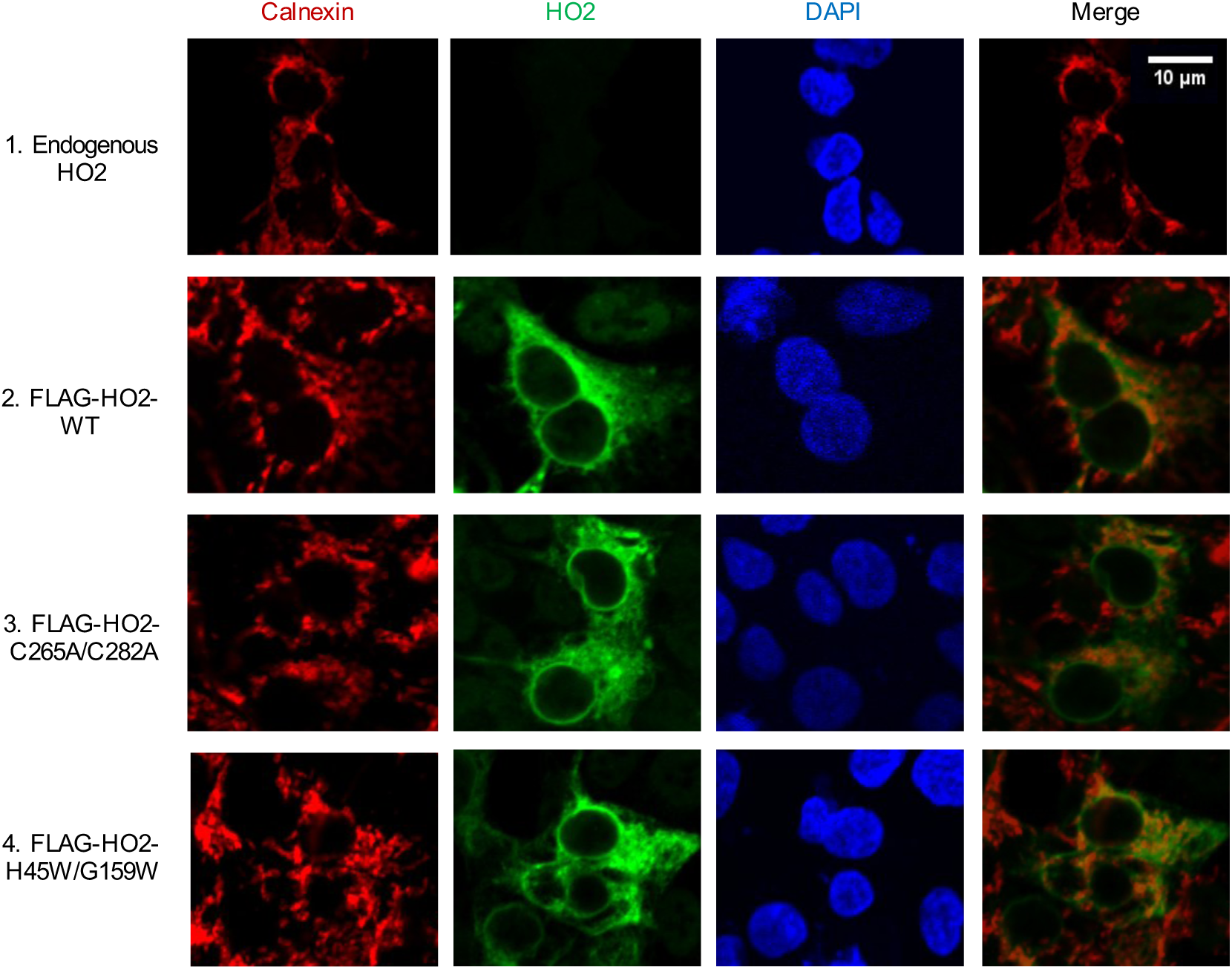
HO2 and its variants localize to the ER. HEK293 cells cultured on coverslips were transfected with transfection reagent only (*Endogenous HO2*) or with corresponding expression vectors for FLAG-tagged HO2 and its variants as indicated. Endogenous HO2 was stained with anti-HO2 antibody, FLAG-tagged HO2 was probed with anti-FLAG antibody. Anti-calnexin antibody was used to visualize calnexin as a marker for ER and DAPI was used to show the position of nuclei. “Merge” is the merged images of Calnexin channel and HO2 channel. All images are in the same scale as indicated with the scale bar.

### Unlike HO1, HO2 is not a substrate of the ERAD pathway

HO1 has been demonstrated to be an ERAD substrate, tagged by poly-ubiquitin and retro-translocated to the proteasome for degradation (12). Based on the high similarity of HO1 and HO2 in structure, catalytic activity and localization, we predicted that HO2 is also degraded through the ubiquitin-proteasome pathway. However, in the presence or absence of CHX, incubation with the proteasome inhibitor MG-132 did not increase levels of either endogenous (Fig. S6) or transiently overexpressed HO2 (Fig. 6A-6B). Furthermore, FLAG-HO2 was immunoprecipitated from HEK293 cell extracts treated with or without MG-132, then detected with a poly-ubiquitin specific antibody. Ubiquitinated HO2 was not detected in the immunoprecipitants, nor did we observe FLAG-HO2 accumulation with MG-132 treatment (Fig. 6C). These combined results indicate that the ubiquitin-proteasome machinery is not required for HO2 degradation making HO2 distinct from other HRM-containing proteins and HO1 in their degradation pathways (18,21,24,36).

**Figure 6.**
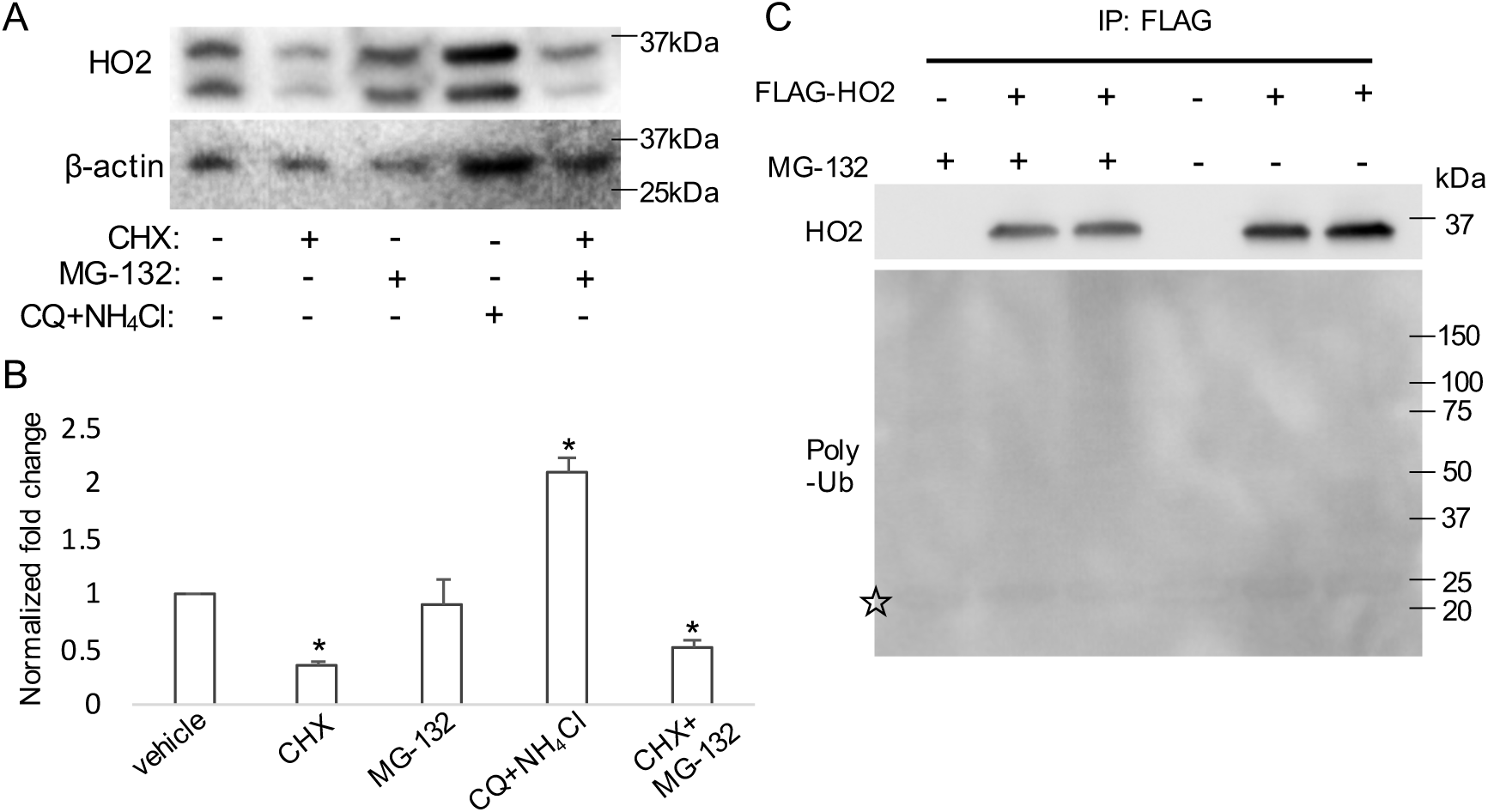
HO2 degradation is not affected by inhibitor of the proteasome. (***A***) HEK293 cells expressing wild-type HO2 were treated with or without CHX (35 μg/ml) in combination with or without MG-132 (20 μM) or chloroquine (CQ)/NH_4_Cl (50 μM and 5 mM, respectively) as indicated for 12h before collection. A representative western blot of samples probed with anti-HO2 antibody (*top panel*), or anti-actin antibody as a loading control (*bottom panel*) is shown. (***B***) The intensities of the bands in (*A*) were quantified by densitometry, and the mean intensities ± S.D. of at least three independent experiments were plotted. *p<0.05 vs vehicle. (***C***) HEK293 cells were transfected pQCXIP empty vector or corresponding expression vector for FLAG-HO2 in pQCXIP, followed by treatment with or without 20 μM of MG-132 as indicated for 12h. Cell lysates were prepared and subjected to immunoprecipitation with an anti-FLAG antibody. The immunoprecipitants were then analyzed by western blot with antibodies against HO2 (*top panel*) or poly-ubiquitin (*bottom panel*). Non-specific band is marked with ☆.

### HO2 is degraded by lysosomes through chaperone-mediated autophagy (CMA)

While MG-132 did not enhance the steady-state level of HO2, treatment with a combination of the lysosomal inhibitors chloroquine (CQ) and NH_4_Cl led to a two-fold increase, suggesting that HO2 is degraded through the autophagy-lysosomal pathway (Fig. 6A). Then we assessed cellular HO2 level in the presence and absence of CHX and each of three different lysosomal inhibitors, bafilomycin (BAF), CQ, and NH_4_Cl. Each inhibitor significantly attenuated HO2 degradation (Fig. 7A), strongly suggesting that HO2 is degraded by the lysosome.

**Figure 7.**
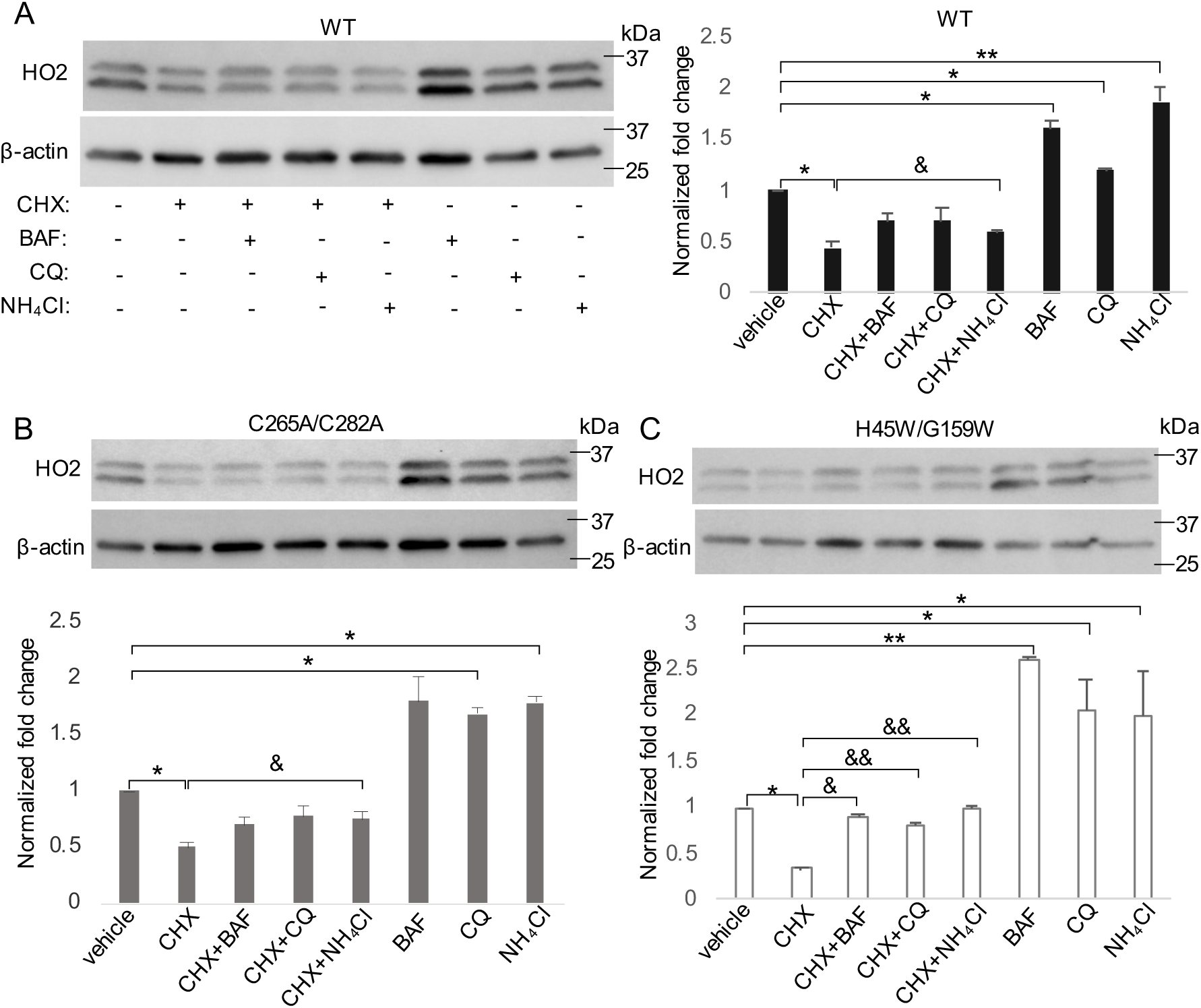
HO2 degradation is blocked by lysosomal inhibitors. (***A-C***) HEK293 cells transfected with expression vector for WT-HO2 (*A*), C265A/C282A mutant (*B*) or H45W/G159W mutant (*C*) treated with or without CHX (35 μg/ml) in combination with or without BAF (10 nM), CQ (50 μM) or NH_4_Cl (10 mM) as indicated for 12h before collection. HO2 protein levels were quantified by western blot analysis and densitometry. Representative western blots of WT, C265A/C282A and H45W/G159W are shown in A (*left panel*), B (*upper panel*), and C (*upper panel*). The order of samples in (*B*) and (*C*) are the same as shown in (*A*). Data are presented as the mean intensities ± S.D. of at least three independent experiments. *p<0.05, **p<0.01 vs vehicle treated group. &p<0.05, &&p<0.01 vs CHX treated group.

We next asked whether heme binding to the catalytic site or to the HRMs impact the HO2 degradation pathway. HEK293 cells expressing C265A/C282A or H45W/G159W variants were subject to the protein degradation assay with or without the aforementioned three lysosomal inhibitors. Each inhibitor blocked lysosomal degradation, leading to protein accumulation of both variants (Fig. 7B-7C). These results indicate that heme binding to the catalytic site controls the rate of HO2 protein degradation but not the degradation pathway. Furthermore, comparison of the normalized fold-changes in HO2 levels between wild-type (Fig. 7A) and the H45W/G159W variant (Fig. 7C) shows that, in the presence of lysosomal inhibitors, accumulation of the H45W/G159W variant is significantly greater than that of the wild-type protein. This result validates that when heme is absent from the catalytic site, HO2 is more vulnerable to lysosomal protein degradation.

The three lysosome inhibitors described above repress both macroautophagy-and CMA-mediated lysosomal degradation. Therefore, to further assess which mode of lysosomal degradation HO2 is subject to, we tested steady-state levels of HO2 in the presence of 3-methyladenine (3-MA), an inhibitor that only blocks macroautophagy (37), or 6-aminonicotinamide (6-AN), which specifically activates CMA (38). The addition of 3-MA did not affect steady-state level of wild-type HO2, while treatment with 6-AN caused a significant decrease in HO2 level (Fig. 8A-8B). Thus, we conclude that CMA mediates HO2 protein degradation.

**Figure 8.**
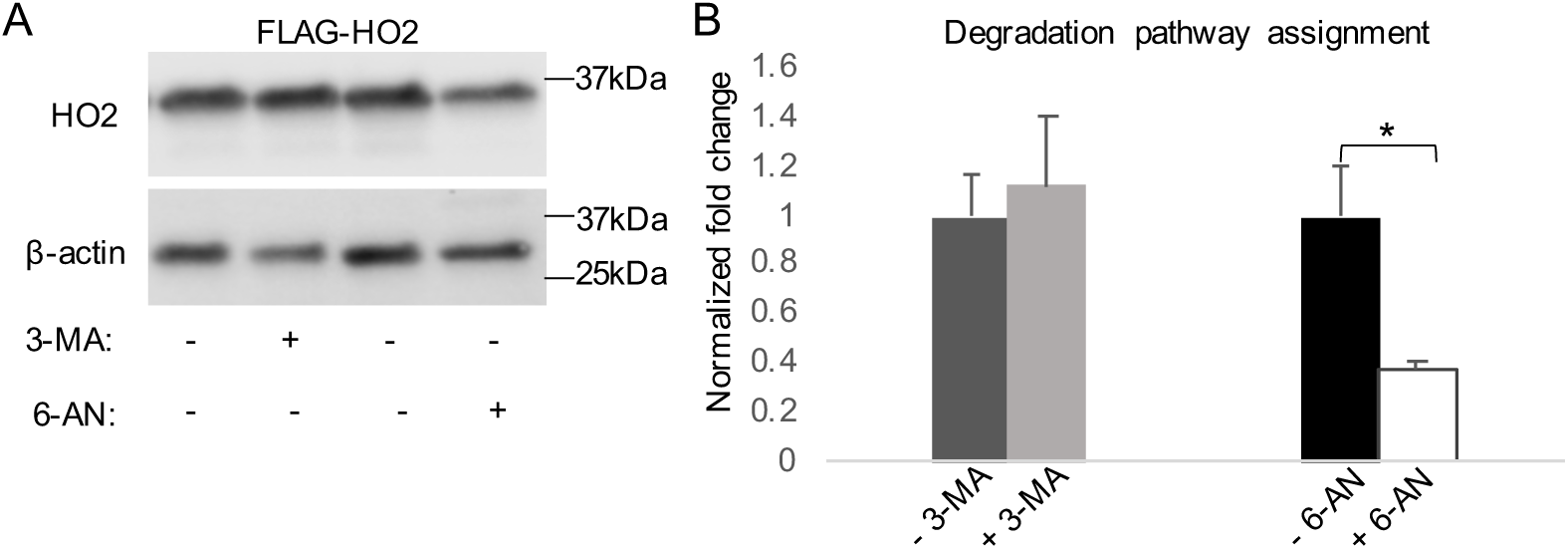
HO2 is degraded by chaperone-mediated autophagy (CMA). (***A)*** HEK293 cells expressing FLAG-HO2-WT were treated with 5 mM of 3-MA for 12h or 50 mM of 6-AN for 24h before collection. A representative western blot of samples probed with anti-HO2 antibody (*top panel*) or anti-actin antibody as a loading control (*bottom panel*) is shown. (***B***) HO2 protein levels were quantified by densitometry. Data are presented as the mean intensities ± S.D. of at least three independent experiments. *p<0.05 vs cells received vehicle DMSO (−6-AN).

## DISCUSSION

It has been unclear how HO2 is regulated to maintain cellular heme homeostasis. In this study, we investigated how the cellular protein level of HO2 is controlled post-translationally by heme. The possession of two additional heme binding sites at its HRMs is the most intriguing feature that distinguishes HO2 from HO1. The most commonly reported HRM function is to promote target protein ubiquitination and proteasomal degradation upon heme binding (18). However, as shown here, heme binding to the HRMs of HO2 does not affect either its steady-state expression level or its protein degradation rate. Furthermore, addition of excess cellular heme does not promote HO2 degradation. Although these properties contrast with those of other HRM-containing proteins, they are consistent with HO2’s role as a heme detoxification enzyme— it would be counterproductive for heme to reduce HO2 level and hence its heme metabolic activity. In light of HRM’s function in HO2, we recently proposed a heme transfer model in which HO2 readily transfers HRM-bound Fe^3+^-heme to the catalytic site for degradation to facilitate turnover and equilibrates heme between the HRMs and the catalytic core to maintain heme homeostasis through heme-dependent conformational changes in the protein (30). This model defines a novel function for the HRM, while maintaining consistency with a conformational change model proposed for another HRM-containing protein, Bach1, in which heme induces local conformational changes that alter interactions with its binding partners (39). It also integrates the thermodynamic properties of the these three heme binding sites in HO2, with the catalytic site exhibiting the highest affinity for heme and the HRMs binding it more weakly (15).

The most surprising finding described here, based on our characterization of several catalytic site variants of HO2 (Fig. 1A), is a previously unrecognized regulatory function of the active site beyond catalysis. Based on our comparison of the steady-state expression levels and degradation rates of these variants versus the wild-type protein, we propose that, when heme is absent from the catalytic site, likely under the condition of heme deficiency, HO2 is destabilized and more subject to degradation (Fig. 3). This proposal agrees with HO2’s function in maintenance of heme homeostasis, since when the cellular heme concentration decreases below its normal physiological level, less heme oxygenase activity is required. Consistently, it has been described that in neuronal cells where HO2 is highly abundant, heme deficiency leads to undesired iron accumulation (40), which can potentially be attenuated by decreasing cellular HO2 level and activity. Given that HO1 is upregulated in response to excess cellular heme, we propose that HO2 helps to restore heme homeostasis by decreasing its own protein level in response to cellular heme deficiency. These proposals offer a rationale for why cells express two HO isoforms: HO1, whose expression levels are highly responsive to excess heme, which accumulates due to cellular injury and stress; whereas HO2, though possessing similar catalytic activity as HO1, exhibits a dual role of heme degradation under physiological conditions and of restoring the regulatory heme pool under conditions of heme deficiency. A similar situation, where two heme oxygenase isoforms are differentially regulated by heme, has been observed in the Isd heme-iron acquisition system of *Staphylococcus aureus* (41). The Isd system contains two heme oxygenases—IsdI and IsdG, which share high structural and sequence homology and possess similar catalytic activities; however, they are differentially regulated by heme and iron (42-44). While the stability and degradation of IsdI is not affected by heme, IsdG is destabilized when heme is absent from the catalytic site (42); furthermore, a catalytically inactive mutant of IsdG is as stable as the wild-type protein in the presence of heme (41), similar to what we describe here for wild-type HO2 and the inactive H45A variant.

It is very interesting that it is the absence of heme binding to the active site (mimicked by H45W/G159W), not the absence of catalytic activity (mimicked by H45A) that triggers HO2 protein degradation. This finding clearly shows that the degradation signal is not triggered by changes in the levels of the products (CO, Fe, biliverdin) of heme degradation. Our recent hydrogen-deuterium exchange mass spectrometry (HDX-MS) studies demonstrated that in H45W/G159W variant, the tryptophan mutations themselves led to minimal to no structural perturbations and that the only observed differences in HDX-MS related to heme binding to the catalytic core were specific to the distal and proximal helices, which form the heme binding pocket (45). Furthermore, the circular dichroism spectra of wild-type HO2 and the H45W/G159W variant were nearly indistinguishable (30). In a search for potential signals for HO2 degradation in the absence of heme, we identified a KFERQ-like motif, which has been classified as a recognition motif by CMA. In the process of CMA, the KFERQ-like motif in target protein is exposed and bound by the heat shock cognate protein of 70 kDa (HSC70), followed by unfolding and translocation of the protein substrate across the lysosomal membrane by lysosome-associated membrane protein type 2A (LAMP 2A) (46). In HO2, one KFERQ-like motif, 52[QFVKD]57, absent in HO1, is found at the proximal helix, which harbors the heme ligand His45 at the HO2 catalytic site. According to our HDX-MS analysis, compared with an apo-active site, decreased deuterium incorporation at the proximal helix was observed when heme is bound at the catalytic site (45). This heme-dependent protection of the KFERQ-like motif could explain how the apo active site of HO2 is differentially recognized by the lysosomal degradation machinery, leading to more rapid protein degradation. Another potential signal for HO2 lysosomal degradation is a di-leucine motif (32[DLSELL]37), which usually serves as sorting sequence for target protein delivery to the lysosomes (47). This motif, not conserved in HO1, also locates at the proximal helix and undergoes similar heme-dependent protection as the aforementioned KFERQ-like motif. Future studies of the di-leucine motif and KFERQ-like motif will be important in better understanding the mechanism of HO2 degradation.

It is also intriguing that HO1 and HO2 are targeted by different protein degradation machinery. CMA has recently been shown to be involved in the execution of ferroptosis, a necrosis related to iron overload (48). It is possible that defects in heme biosynthesis that leads to cellular iron overload induces ferroptosis (40). Therefore, CMA is already activated when a rapid HO2 degradation is required, providing a convenient pathway. Future studies will focus on whether heme deficiency activates CMA, therefore enhancing HO2 degradation.

Even though stabilization of an enzyme by substrate binding is not an uncommon feature, in the case of HO2 it is surprising because the stabilization occurs at concentrations two-orders-of magnitude lower than the Michaelis constant (K_m)_. Since HO2 degradation is modulated only by heme binding but not by heme metabolism at the catalytic site, it is worth considering the relative K_m_ (0.40 µM) (49) and K_d_ values for heme (3.6 nM for the catalytic site) (28) in relation to the total (∼3.2 µM) and regulatory (20–340 nM) heme levels found in mammalian cells (29). The agreement of the K_d_ value with the regulatory heme pool suggests that the degradation signal of HO2 is related to changes in the regulatory, not the total, heme pool. Consistently, HO2 overexpression in mammalian cells does not alter total heme level but only shifts the regulatory heme pool (personal communication with Prof. Amit R. Reddi). Thus, degradation of the housekeeping heme detoxification enzyme HO2 is likely to be linked to the larger network of regulatory heme trafficking within the cell. The detailed mechanism of how this heme-dependent HO2 degradation helps to restore cellular heme homeostasis under heme deficiency is open for investigation.

## EXPERIMENTAL PROCEDURES

### Materials

Cycloheximide (CHX), MG-132, bafilomycin (BAF), 3-methyladenine (3-MA), 6-aminonicotinamide (6-AN) and poly-lysine were purchased from Sigma-Aldrich. Chloroquine and NH_4_Cl were from Fisher. Fetal bovine serum (FBS), Dulbecco’s Modified Eagle medium (DMEM), Opti-MEM, Lipofectamine LTX&PLUS reagent and a mounting reagent with DAPI were from Invitrogen. A rabbit polyclonal heme oxygenase 2 antibody (ab90515) was purchased from Abcam. A mouse monoclonal M2-Flag antibody (F1804), a rabbit polyclonal β-actin antibody (A2066) and a rabbit monoclonal Flag antibody (F7425) were from Sigma-Aldrich. A mouse monoclonal Calnexin antibody (MA3-027) was from Invitrogen. A rabbit polyclonal Ub Antibody (FL-76) (sc-9133) was from Santa Cruz Biotechnology. An anti-Flag M2 magnetic bead (M8823) was purchased from Sigma. Goat-anti-mouse-Alexa 568 antibody and goat-anti-rabbit-Alexa 488 antibody (Molecular Probes) were generously provided by Dr. Phyllis I. Hanson (University of Michigan).

### Plasmid constructs and mutagenesis

Full-length human HO2 cDNA was subcloned into a pcDNA3.1 (+) vector using Hind III and Xho I restriction sites (14). The Flag-tagged WT-HO2 construct was generously provided by Dr. Stephen P. Goff (Columbia University). HO2 point mutations were generated by either site-directed mutagenesis with the Quick-Change protocol (Agilent Technologies) or the Q5 site-directed mutagenesis kit (NEB E0554S).

### Cell culture conditions

HEK293 cells were obtained from ATCC and cultured in basal media (1x DMEM containing 10% FBS and 1% penicillin-streptomycin). Cells were reseeded at 24-48 h prior to transfection (with lipofectamine LTX & PLUS reagent in Opti-mem) and various treatments. For transient transfection experiments, HEK293 cells were transfected with plasmids expressing HO2 or its variants using the Lipofectamine LTX transfection reagent according to the manufacturer’s instructions. After 24 h, cells were subjected to various treatments as indicated (24).

### Protein degradation assay and degradation pathway assignment

Protein stability and degradation rate was determined by the CHX degradation assay, as described before with slight modifications (24). Briefly, for the protein half-life assessment, HEK293 cells were transfected with different constructs for expressing HO2 and its variants. 24h after the transfection, cells were treated with 35 μg/ml of CHX for the indicated time before collection. For degradation pathway assignment, 24h after transfection, cells were treated with indicated inhibitors (MG-132 at 20 μM, BAF at 10 nM, chloroquine at 50 μM or NH_4_Cl at 10 mM) for 12h before harvesting (12). For studying the involvement of different types of autophagy in HO2 degradation, HEK293 cells were transfected to express wild type-HO2. 24h after transfection, cells were treated with 3-MA or vehicle media for 12h before harvesting to probe the effect of macroautophagy on HO2 degradation. For testing chaperone-mediated autophagy, cells were treated with 6-AN or vehicle DMSO for 24h before collection (50). Cell pellets were lysed with RIPA buffer (Thermo scientific) containing 2x protease inhibitor cocktail (Roche); centrifuged at 12,000 rpm for 10 min at 4 °C to get rid of cell debris. The supernatant was extracted to a clean tube and protein concentrations were determined by bicinchoninic acid protein assay (Pierce BCA protein assay kit). Samples were analyzed by western blot.

### Protein assays, SDS-PAGE, and western blotting analysis

All protein concentrations were analyzed by the BCA assay before being further processed. Same amount of total protein from each sample was loaded onto the SDS gel. SDS-PAGE was performed with Laemmli buffers (51), 4–20% gradient gels (Bio-Rad), and protein bands were visualized with Coomassie Brilliant Blue (Bio-Rad) solution or transferred to PVDF membranes (Bio-Rad). Transfer was performed in transfer buffer (25 mM Tris, 192 mM glycine, and 0.025% SDS) at 4 °C, 90 V for 1.5 h. Membranes were blocked in 5% blotting-grade blocker (Bio-Rad) in 1xTBST (20 mM Tris, pH 7.4, 200 mM NaCl and 0.1% Tween 20) prior to incubation with primary antibodies overnight at 4 °C. All primary antibodies were diluted in TBST containing 0.1% sodium azide and 5% BSA as follows: HO2 (1:1,000), β-actin (1:2,000), FLAG-mouse monoclonal (1:1,000) and poly-Ub (1:1,000). After removal of primary antibody, membranes were washed with 1x TBST for 5x 5min and incubated with HRP-conjugated secondary antibodies against the species that primary antibodies were produced in. All secondary antibody was diluted with 5% blotting-grade blocker in TBST with a dilution range from 1:50,000 to 1:30,000. Membranes were washed 5×5min with TBST again before developed with SuperSignal West Femto enhanced chemiluminescent substrate for HRP (Thermo scientific). Immunoreactive bands were detected using a ChemiDoc MP Imaging System from BIO-RAD.

### Protein expression, purification, and analysis

A truncated, soluble form of HO2 spanning residues 1-248 (the core of the protein) was expressed from the pET28a vector in BL21(DE) cells (Life Technologies) and purified by Ni-NTA– agarose affinity chromatography (Qiagen) as described previously (15). The N-terminal six-His tag was removed by treatment with thrombin prior to use (15). The G163H (30) and G163D variants were constructed using the Q5 site-directed mutagenesis kit (New England Biolabs) to introduce the mutation, and the H45A variant was constructed by site-directed mutagenesis with the Quik Change protocol (Agilent Technologies). All variants were purified as described for the wild type protein. All proteins were extensively dialyzed against 50 mM Tris buffer pH 8.0/50 mM KCl prior to use, and protein concentrations were determined by Bradford assay.

A heme stock was freshly prepared in 50 mM Tris buffer pH 8.0/50 mM KCl with 15% dimethyl sulfoxide and 0.1 M NaOH. The stock was passed through a 22 μm filter to remove insoluble matter, and the concentration was determined by using an ε_385_ of 58.4 mM^−1^ cm^−1^ (52). Wild type HO2 and variants (3 mM) were incubated with 2.5 mM heme for 1 hour, and then absorbance spectra were measured at 20 °C in 50 mM Tris buffer pH 8.0/50 mM KCl on a Shimadzu UV-2600 UV–vis spectrophotometer. The heme-bound proteins were subsequently mixed with apo-H64Y/V68F myoglobin (30 mM), and the absorbance at 600 nm over time was recorded. Rates were determined by fits of the data to a double exponential equation using GraphPad Prism 8. H64Y/V68F-myoglobin (green heme) was purified and converted to the apo-form as described previously (28) using an expression vector generously supplied by J. Olson (Rice University, Houston, TX).

Steady state activity of HO2 was measured as described previously (53) with minor modifications. Briefly, a 200 mL reaction containing 0.1 mM HO2, 15 mM heme, 1 mM biliverdin reductase, 0.25 mg/mL bovine serum albumin, 10 units catalase, and 10 mM cytochrome P450 reductase in 50 mM Tris pH 8.0, 50 mM KCl was incubated for 2 minutes at 37 °C. The reaction was initiated by the addition of 1 mL of 100 mM NADPH. Activity was monitored by following the increase in absorbance at 468 nm due to bilirubin formation. Using a difference extinction coefficient of 43.5 mM^-1^ cm^-1^, activity was calculated as nmol bilirubin formed per minute per mg HO2. Cytochrome P450 reductase and biliverdin reductase were purified as described previously (53).

### HO2 immunoprecipitation

HEK293 cells were transfected to express FLAG-tagged HO2 or its variants for 24h before harvesting for immunoprecipitation assay (54). Cells were lysed in CelLytic M Cell Lysis Reagent (Sigma, C2978) on ice for 20 min with mild agitation. The lysate was centrifuged for 12 min at 12000 rpm and 4°C to clear out cell debris. Immunoprecipitation was performed as the manufacturer instructed. Briefly, the supernatant was mixed with ANTI-FLAG M2 magnetic bead (M8823) and the mixture was incubated at 4°C for 4 h. After removing the flow through, the magnetic bead was washed with lysis reagent once and TBST twice, 5 min for each wash at 4 °C. Protein complexes were eluted in SDS-PAGE sample buffer (125mM Tris-HCl, pH6.8, 4% SDS, 20% glycerol and 0.004% bromophenol blue) with boiling for 10min. Eluted samples were subjected to electrophoresis and western blot detection.

### Cellular heme concentration modulation

Modulation of intracellular heme concentration was performed as described before (17) with modifications. Briefly, the cells cultured with regular media were collected from confluent master plates, and reseeded into 35mm dishes with 1:10 dilution in fresh basal media on day 0. On day 2, media was replaced with heme-depleted (HD) media (1x DMEM + 10% heme-depleted FBS) and cells were transfected with 1μg of plasmids for HO2 or its variants expression in Opti-mem with 0.5 mM succinylacetone (SA). On day 4, the media was replaced with fresh HD media containing 0.5 mM succinyl acetone, and supplemented with 0, 10 or 50 μM hemin. After 24h of media exchange, cells were harvested for further analysis.

### Oxalic acid assay for total heme content

After modulating intracellular heme concentration, the heme level in soluble cell extracts was determined by iron extraction with a supersaturated oxalic acid solution and measurement of the resulting protoporphyrin fluorescence as described previously (24,55) with slight modifications. Briefly, cells were lysed in CelLytic M Cell Lysis Reagent (Sigma, C2978) containing 2x protease inhibitor cocktail (Roche); protein concentrations were determined by BCA assay. Cell extracts were diluted at 1:20 in supersaturated oxalic acid in sealed glass cryo vials, mixed, and then heated at 121 °C for 30 min in an autoclave. A parallel set of samples was left at room temperature. The heated samples were cooled down and transferred to a black 96-well plate. The fluorescence emission spectrum of protoporphyrin was recorded from 550–700 nm with excitation at 400 nm in a Tecan Safire microplate reader. A standard curve was generated with hemin prepared in DMSO. Heme concentration (nmol per mg protein) was quantified after subtraction of background emission from non-heme protoporphyrin in the unheated samples from the intensity of the heated samples at 662 nm.

### Immunofluorescence microscopy (IF)

HEK293 cells were cultured on poly-lysine coated coverslips for 24h and transfected with plasmids expressing FLAG-tagged HO2 and its variants. 24h after transfection, cells were washed with phosphate buffered saline (PBS) twice and fixed with 4% paraformaldehyde for 10min at room temperature. IF samples were then washed with PBS twice more before being permeabilized with 1x PBS containing 0.2% Triton X-100 for 10min then washed with PBS containing 0.1% Tween 20 (PBST) for 3x 5min. Permeabilized samples were transferred to a new 12-well plate and blocked with blocking buffer (PBST + 0.3% BSA) for 1h. Wash coverslips thoroughly with PBST for 3x 5min and incubated with primary antibody diluted in PBST containing 1% BSA for 1h. After removing primary antibody, coverslips were washed again with PBST for 3x 5min and incubated with secondary antibody for 1h. Coverslips were washed with PBST before mounted onto microscope slides. Sample images were acquired on an Olympus IX81 microscope with a Yokogawa CSU-W1 spinning disk confocal scanner unit, an Olympus PlanApo 60x 1.42 NA objective, and a Hamamatsu ImagEMX2-1K EM-CCD digital camera. DAPI and secondary antibodies conjugated to Alexa Fluor 488 and Alexa Fluor 568 were excited with 405 nm (Omicron), 488 nm (Omicron), and 561 nm (Coherent) solid state lasers paired with Chroma ET460/50m, ET525/50m, and ET605/52m emission filters, respectively. Images were acquired with MetaMorph 7.10.2.240 and prepared in FIJI software. Primary antibodies were applied as follow: HO2 (1:200), Calnexin (1:200) and FLAG (1:100). Secondary antibodies were applied as follow: Goat anti mouse-Alexa 568 (1:1,000) and Goat anti rabbit-Alexa 488 (1:1,000). The mounting reagent contains DAPI.

### Statistical analysis

Data values of at least three independent experiments were analyzed by a two-tailed unpaired student’s t-test. Significance was established at p < 0.05. Error bars represent +/-SD.

## ABBREVIATIONS

The abbreviations used are:

6-AN: 6-aminonicotinamide
ALAS1: d-aminolevulinic acid synthase 1
BAF: bafilomycin
CHX: cycloheximide
CMA: chaperone-mediated autophagy
CQ: chloroquine
CVD: cardiovascular disease
DMSO: dimethyl sulfoxide
ERAD: endoplasmic reticulum-associated degradation
HD: heme-depleted
HDX-MS: hydrogen-deuterium exchange mass spectrometry
HO1: heme oxygenase-1
HO2: heme oxygenase-2
HRM: heme regulatory motif
HSC70: heat shock cognate protein of 70 kDa
LAMP 2A: lysosomal-associated membrane protein type 2A
3-MA: 3-methyladenine
Ni-NTA: Ni-nitrilotriacetic acid
ROS: reactive oxygen species
SA: succinylacetone.

## ACKNOWLEDGMENTS

We thank Prof. Amit R. Reddi, and Dr. David A. Hanna for critical reading of the manuscript; Prof. Phyllis I. Hanson and Dr. Kevin Bohannon for assistance with microscopy; and Prof. Stephen P. Goff for providing the FLAG-tag expression plasmid. The content is solely the responsibility of the authors and does not necessarily represent the official views of the National Institutes of Health.

